# HBx enhances CPAP expression via interacting with CREB to promote hepatocarcinogenesis in HBV-associated HCC

**DOI:** 10.1101/423194

**Authors:** Chia-Jui Yen, Shu-Ting Yang, Ruo-Yu Chen, Wenya Huang, Kazuaki Chayama, Ming-Hao Lee, Shiang-Jie Yang, Yu-Wei Hsiao, Ju-Ming Wang, Yih-Jyh Lin, Liang-Yi Hung

**Author notes:** These authors contributed equally to this work (Chia-Jui Yen, Shu-Ting Yang). Contact informations: Liang-Yi Hung and Yih-Jyh Lin, Department of Biotechnology and Bioindustry Sciences, College of Bioscience and Biotechnology, National Cheng Kung University, Tainan 70101, Taiwan; and Division of General and Transplantation Surgery, Department of Surgery, National Cheng Kung University Hospital, College of Medicine, National Cheng Kung University, Tainan 70101, Taiwan.

## Abstract

Hepatitis B virus (HBV) encoded non-structure protein X (HBx) can promote cell proliferation, migration, and anti-apoptosis via activating several transcription factors and increasing their downstream gene expression in HBV-infected liver cells. Our previous report suggested that centrosomal P4.1-associated protein (CPAP) is required for HBx-mediated NF-κB activation. Here, we found that, upon HBV infection, overexpressed HBx can transcriptionally up-regulate *CPAP* via interacting with CREB. CPAP can directly interact with HBx to promote HBx-mediated cell proliferation and migration; and SUMO modification of CPAP is involved in interacting with HBx. Interestingly, CPAP can increase the HBx protein stability in an NF-κB-dependent manner; and overexpressed CPAP and HBx is positively correlated with the activation status of NF-κB in HCC. Increased expression of *CREB* and *CPAP* mRNAs exists in the high-risk group with a lower survival rate in hepatocellular carcinoma (HCC). These results suggest that the reciprocal regulation between CPAP and HBx may provide a microenvironment to facilitate HCC development via enhancing NF-κB activation, inflammatory cytokine production, and cancer maligancies. The findings of this study not only shed light on the role of CPAP in HBV-associated HCC, but also provide CPAP as a potential target for HBV-related HCC therapy.

**Author Summary:** In this study, we address a novel molecular mechanism for the collaboration between overexpressed HBx and CPAP in promoting hepatocarcinogenesis in HBV-associated HCC. Upon HBV infection, HBx is overexpressed and interacts with CREB to transcriptionally activate CPAP; the HBx/CPAP interaction promotes hepatocarcinogenesis. Clinical analysis found that co-overexpressed CPAP and CREB exist in the high-risk group with a lower survival rate in HCC. Additionally, overexpressed CPAP contributes to HBx protein stability in a NF-κB-dependent pathway. Our study provides a potential translational application in targeting CREB-CPAP axis in HBV-associated HCC.

## Introduction

HBV infection is a major etiological factor in acute and chronic hepatitis and enhances the development of liver diseases such as cirrhosis and HCC. Among the different HBV proteins encoded by HBV genome, the X protein of HBV (HBx) plays a critical role in HBV-associated HCC development possibly through triggering specific oncogenic pathways and causing an accumulation of genetic and epigenetic alterations in regulatory genes(1-4).

HBx is a multifunctional protein that modulates the expression of various cellular and viral genes involving in cell survival, cell cycle progression, DNA repair, invasion, protein degradation and regulates several signaling pathways such as Ras/Raf/MAPK, PI3K/Akt, NF-κB or JNK(5-10). Additionally, functions of HBx on cytoplasmic signal transduction cascades and transcriptional activation imply that HBx is both a cytoplasmic and nuclear protein(11). It is known that HBx serves as a powerful transcriptional activator that up-regulates several transcription factors including NF-κB, AP-1, and even the HBV genome(12, 13). However, HBx does not bind DNA directly. HBx affects transcription by interacting with several transcription factors, including DNA-binding factors and complexes of transcriptional machinery(14). For example, HBx interacts with TATA box-binding protein (TBP), RNA polymerase II subunit 5 (RPB5), and transcription factor IIB (TFIIB) to regulate RNA polymerase transcription(15-17). In the nucleus, HBx associates with C-terminal binding protein (CBP/p300) and binds to the CREB-targeting site of the promoters of *IL-8* and proliferating cell nuclear antigen (*PCNA*)(18, 19). Recently, an *HBx* transgenic mouse model showed a high incidence of liver tumor formation without fibrosis in 90% of cases and has been widely used as an animal model for studying the detailed mechanisms of chronic HBV infection in HCC development(20, 21). Although the role of HBx in the pathogenesis of HCC is well understood, the mechanism by which HBx regulates the gene expression network is not fully clear.

Previously, we showed that the expression of CPAP in HBV-associated HCC correlates with a poor prognosis(22). CPAP has been reported to be part of the γ-tubulin complex, which is associated with γ-tubulin in both the centrosomal and cytosolic fractions throughout the cell cycle, and plays an essential role in microtubule nucleation and procentriole elongation(23-25). Interestingly, CPAP also regulates cell apoptosis and the growth of neural precursor cells(26, 27). There are three nuclear localization signals and two nuclear export signals within the CPAP polypeptide(28), indicating CPAP can shuttle between the nucleus and cytoplasm. Furthermore, CPAP has been shown to act as a transcriptional coactivator of STAT5 and NF-κB(28, 29). TNF-α-induced SUMO modification of CPAP is required for IKK-mediated NF-κB activation in HCC cell lines and promotes the growth of HCC cells, suggesting that CPAP is critical for the association between NF-κB and inflammation-related diseases, such as HCC(22). In addition, the cooperation of CPAP and HBx in regulating the transcriptional activity of NF-κB, provides evidence that CPAP plays an important role in HBx-mediated HCC development(22). However, the relationship between CPAP and HBx and the physiological role of CPAP in HBV-associated HCC are still unclear.

In this study, we investigated the interaction between CPAP and HBx and determined the functional role of the CPAP-HBx interaction in HBx-mediated hepatocarcinogenesis. HBx transcriptionally increased the expression of *CPAP* via interacting with CREB, and overexpressed CPAP increased the protein stability of HBx in an NF-κB-dependent manner, both of which resulted in an increased activity and target gene expression of NF-κB. The reciprocal regulatory loop between CPAP and HBx at the transcriptional and post-transcriptional levels presents a complex relationship during early and late hepatocarcinogenesis in HBV-related HCC. HCC patients with co-overexpressed *CREB* and *CPAP* mRNAs have a poor prognostic value. Taken together, our results provide strong evidence that CPAP is crucial for HBx-induced HCC development and may offer opportunities to develop mechanism-based therapies.

## Results

### Overexpression of CPAP in HBV-associated HCC

Our previous studies have showed that CPAP expression positively correlates with a poor prognosis in HBV-HCC(22). To further assess the clinical significance of CPAP expression in HBV-HCC, we evaluated the association between CPAP and the major clinicopathological features of 132 HBV-HCCs (Supplementary Table 2). The results showed a significant correlation between high *CPAP* expression levels with the disease-free survival rate, AST, ALT, differentiation, tumor size, and AJCC stage (Table 1).

**Table 1.** Associations between *CPAP* expression, clinical parameters and disease-free survival/ overall survival

| Clinical parameters | N | Disease-free survival (years) |  |  |  | Overall survival (years) |  |  |  |
| --- | --- | --- | --- | --- | --- | --- | --- | --- | --- |
|  |  | Mean | 95% CI |  | p-value | Mean | 95% CI |  | p-value |
| *CPAP* expression |  |  |  |  |  |  |  |  |  |
| < 1.5X | 47 | 2.776 | 2.345 | 3.237 | 0.039* | 3.858 | 3.376 | 4.340 | 0.521 |
| ≥ 1.5X | 85 | 2.644 | 1.815 | 3.472 |  | 7.886 | 7.122 | 8.650 |  |
| Gender |  |  |  |  |  |  |  |  |  |
| Male | 102 | 2.379 | 2.039 | 2.720 | 0.922 | 7.616 | 6.837 | 8.394 | 0.413 |
| Female | 30 | 2.825 | 1.768 | 3.881 |  | 7.983 | 6.941 | 9.025 |  |
| Age (yrs) |  |  |  |  |  |  |  |  |  |
| < 50 | 30 | 2.508 | 1.994 | 3.021 | 0.196 | 4.001 | 3.507 | 4.495 | 0.867 |
| ≥ 50 | 102 | 2.869 | 1.968 | 3.770 |  | 7.736 | 6.987 | 8.486 |  |
| Liver cirrhosis |  |  |  |  |  |  |  |  |  |
| No | 89 | 2.042 | 1.709 | 2.376 | 0.231 | 7.855 | 7.040 | 8.671 | 0.771 |
| Yes | 43 | 3.387 | 2.197 | 4.577 |  | 7.400 | 6.364 | 8.437 |  |
| AST/GOT (U/L) |  |  |  |  |  |  |  |  |  |
| < 52 | 80 | 3.827 | 3.124 | 4.530 | 0.118 | 8.386 | 7.749 | 9.023 | 0.010* |
| ≥ 52 | 52 | 2.189 | 1.658 | 2.720 |  | 6.532 | 5.278 | 7.787 |  |
| ALT/GPT (U/L) |  |  |  |  |  |  |  |  |  |
| < 111 | 120 | 3.083 | 2.156 | 4.010 | 0.074 | 8.030 | 7.423 | 8.637 | 0.045* |
| ≥ 111 | 12 | 1.706 | 0.822 | 2.591 |  | 2.648 | 1.935 | 3.361 |  |
| AFP (ng/ml) |  |  |  |  |  |  |  |  |  |
| < 400 | 103 | 2.954 | 2.048 | 3.860 | 0.929 | 7.825 | 7.091 | 8.559 | 0.510 |
| ≥ 400 | 29 | 2.397 | 1.776 | 3.018 |  | 7.264 | 5.925 | 8.604 |  |
| Differentiation |  |  |  |  |  |  |  |  |  |
| Well | 25 | 2.583 | 1.891 | 3.274 | 0.063 | 7.957 | 7.254 | 8.661 | 0.009* |
| Moderate | 91 | 3.850 | 3.176 | 4.525 |  | 7.754 | 6.950 | 8.559 |  |
| Poor | 16 | 1.207 | 0.764 | 3.900 |  | 3.399 | 2.227 | 4.571 |  |
| Tumor size (cm) |  |  |  |  |  |  |  |  |  |
| < 5 | 87 | 3.536 | 2.455 | 4.617 | 0.000* | 8.346 | 7.822 | 8.870 | 0.000* |
| ≥ 5 | 45 | 1.529 | 1.086 | 1.973 |  | 6.007 | 4.500 | 7.513 |  |
| Tumor number |  |  |  |  |  |  |  |  |  |
| Single | 107 | 3.034 | 2.092 | 3.976 | 0.921 | 7.764 | 7.033 | 8.495 | 0.631 |
| > 1 | 25 | 2.155 | 1.572 | 2.739 |  | 3.227 | 2.735 | 3.718 |  |
| Vascular invasion |  |  |  |  |  |  |  |  |  |
| Absence | 94 | 3.014 | 2.077 | 3.951 | 0.601 | 8.168 | 7.468 | 8.869 | 0.057 |
| Presence | 38 | 2.110 | 1.626 | 2.593 |  | 6.764 | 5.532 | 7.997 |  |
| AJCC Stage |  |  |  |  |  |  |  |  |  |
| I&II | 108 | 3.451 | 2.402 | 4.501 | 0.000* | 8.238 | 7.626 | 8.850 | 0.001* |
| III | 24 | 0.910 | 0.526 | 1.295 |  | 3.074 | 2.281 | 3.867 |  |

### HBx transcriptionally up-regulates CPAP by interacting with CREB

By Western blot analysis, we found that HBx can increase CPAP expression in HCC cells (Figure 1A, left panel). HBx stable expression cell lines Hep3BX or HepG2X also exhibited a higher *CPAP* mRNA expression than Hep3B or HepG2 (Figure 1A, middle panel). The *CPAP* promoter assay further demonstrated that HBx transcriptionally up-regulates the *CPAP* promoter (Figure 1A, right panel, and Supplementary Figure 1). The same result was obtained from HBV-infected HCC or HBV genome-expressing HCC cells(30). Expression of *CPAP* mRNA and protein is increased in HBV-infected (Figure 1B) or HBV genome-expressed HCC (Supplementary Figure 2). To identify the specific transcription factors involved in HBx-enhanced transcriptional activation of *CPAP*, TF SEARCH (http://www.cbrc.jp/research/db/TESEARCH.html) was used to evaluate the *CPAP* promoter, and two potential CREB binding sites (CBSs) were found (Figure 1C, -226 to -215 bp and -86 to -79 bp). The reporter assay showed that *CPAP* promoter activities were significantly decreased when CREB binding site 1 was mutated (M1), whereas a mutation in CREB binding site 2 (M2) only moderately decreased the promoter activity compared with the wild type control (Supplementary Figure 3). Given that HBx has been shown to interact with CREB-binding protein/p300 to mediate CREB-dependent transcription(3, 31), we assumed that HBx might activate the *CPAP* promoter through the interaction with CREB. As anticipated, overexpression of CREB or HBx increased *CPAP* promoter activity, which was further enhanced by co-expression of HBx and CREB (Figure 1C and Supplementary Figure 4A). Overexpression of the CREB dominant-negative (DN) mutant had a smaller effect on *CPAP* promoter activity (Supplementary Figures 4B and 4C); knocked-down expression of *CREB* led to decreased *CPAP* promoter activity (Supplementary Figures 5A and 5B). Either HBx or CREB can induce wild type *CPAP* promoter activity but failed to affect *CPAP*/M1 or M1+2 promoter activities (Figure 1D and Supplementary Figure 4D). Overexpression of HBx retained the ability to activate *CPAP* promoter in *CREB* knocked-down cells (Supplementary Figure 5C), while the CREB expression is still maintained in a low level (Supplementary Figures 5A and 5C).

**Figure 1.**
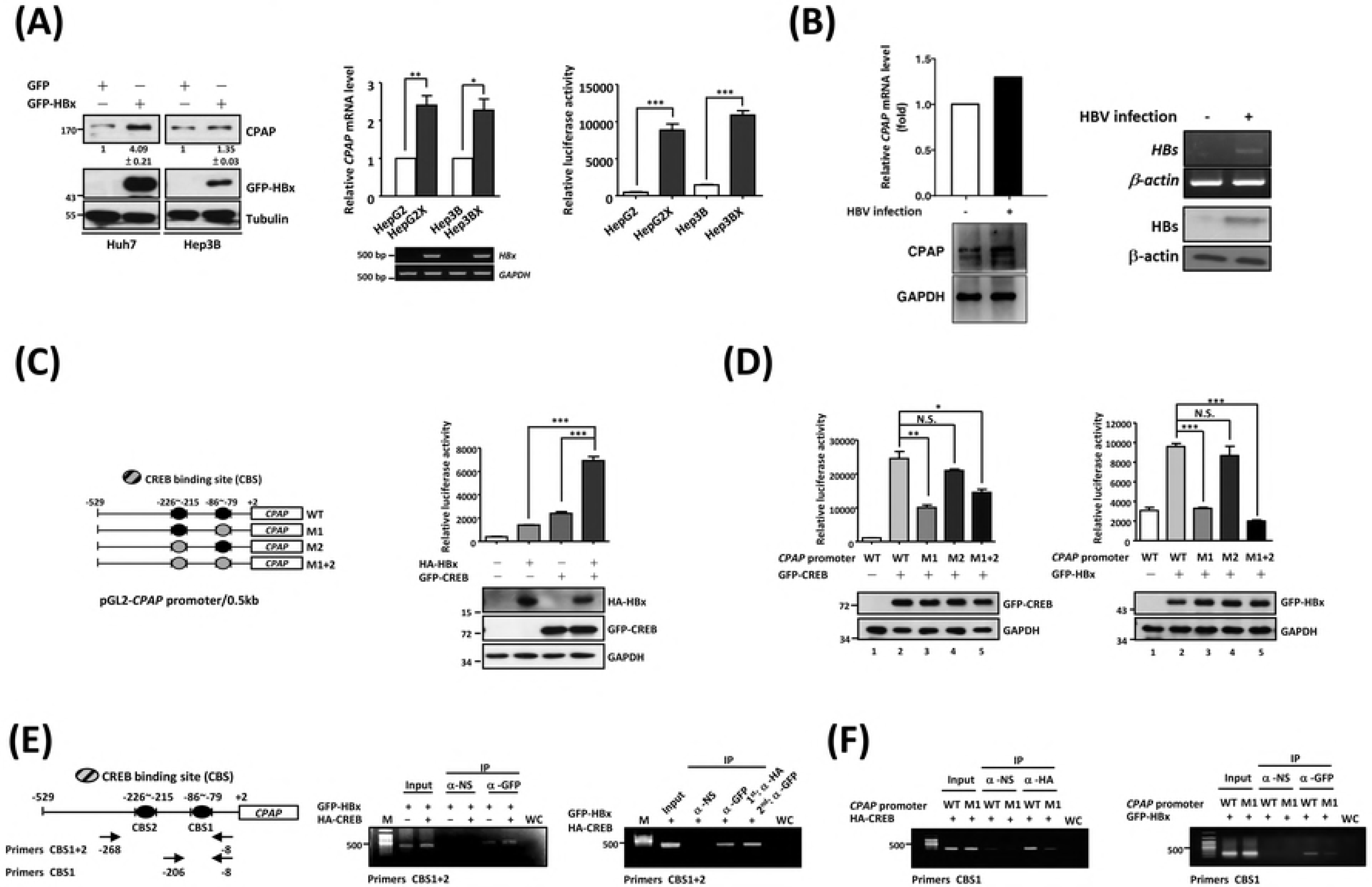
CREB is crucial for HBx to associate with *CPAP* promoter and to enhance the *CPAP* promoter activity. (A) Left) Huh7 or Hep3B cells with GFP or GFP-HBx expression were collected to detect the expression of CPAP and GFP-HBx by Western blot analysis. The mRNA level (middle) and promoter activity (right) of *CPAP* in HepG2, HepG2X, Hep3B and Hep3BX cells were determined by Q-PCR (for *CPAP* mRNA) or RT-PCR (for *HBx* mRNA) and reporter assay, respectively. The expression levels of endogenous CPAP are indicated as ratio (CPAP/Tubulin). The mean ± S.D. was obtained from triplex Q-PCR, and three independent experiments were performed. Error bars of reporter assay represent the mean ± SD of three independent experiments, each performed in triplicate. **(B)** The expressions of CPAP (left) and HBV surface antigen (HBs) (right) were determined by Q-PCR or RT-PCR and Western blot analysis in HBV-infected (+) or no infected (-) HepG2-NTCP-C4 cells(30). **(C)** (Left) A schematic representation of the *CPAP* proximal promoter (pGL2-*CPAP* promoter/0.5 kb) is shown. Two potential CREB binding sites (CBSs) are showed in black ovals (WT), and the promoter with either one or both CBSs mutations is presented by gray color (M1, M2, or M1+M2). (Right) *CPAP* promoter activity was examined in Huh7 cells with GFP-CREB or/and HA-HBx expression by reporter assay. **(D)** pGL2-*CPAP*/WT, M1, M2 or M1+2 was transiently transfected into Huh7 cells with GFP-CREB (left) or GFP-HBx (right), and then, the luciferase activity was analyzed. Error bars represent the mean ± SD of three independent experiments, each performed in triplicate. *, *p* < 0.05; **, *p* < 0.01;***, *p* < 0.001; N.S., no significance. **(E)** (Left) Primers used in chromatin immunoprecipitation (Ch-IP) and re-ChIP assay of the *CPAP* promoter are shown. The ChIP (middle) and re-ChIP (right) assay were performed in GFP-HBx- and HA-CREB-expressing Huh7 cells by anti-GFP, anti-HA antibody and normal serum control. **(F)** pGL2-*CPAP*/WT or M1 and HA-CREB (left) or GFP-HBx (right) were co-transfected into Huh7 cells, and then analyzed by ChIP assay using anti-HA or anti-GFP antibody as described in Fig. 1E. WC: water control.

**Figure 2.**
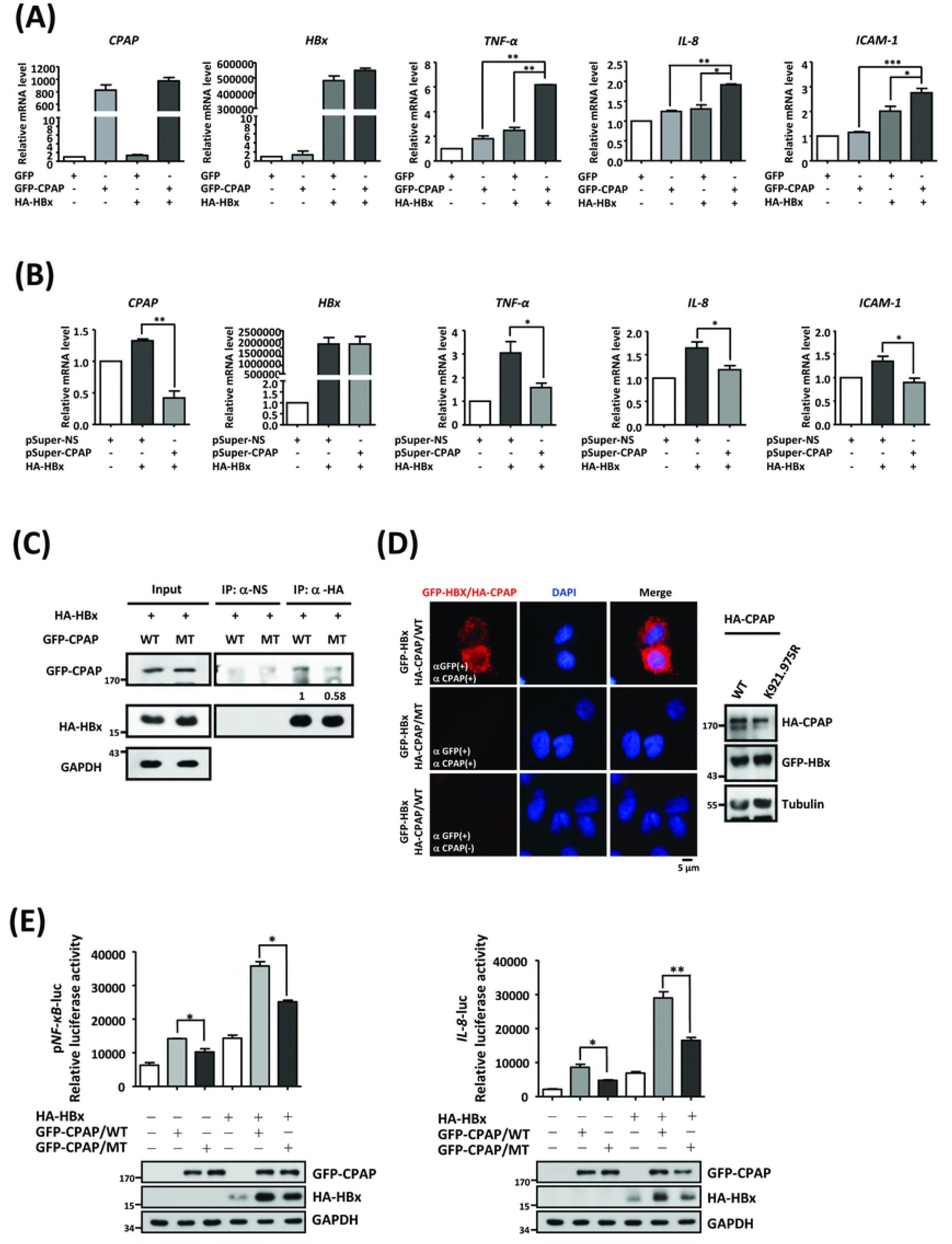
CPAP interacts and cooperates with HBx to enhance NF-κB target genes expression. (A-B) The mRNA level of NF-κB-mediated target genes (*TNF-α, IL-8, ICAM-1*) were detected by Q-PCR in Huh7 cells with co-expressed HA-HBx and GFP-CPAP (A) or knockdown of CPAP (B, pSuper-CPAP). Data present three independent experiments, and each performed in triplicate (*, *p* < 0.05; **, *p* < 0.01;***, *p* < 0.001). **(C)** Immunoprecipitation (IP) assay was performed in total cell lysates from Huh7 with GFP-CPAP/WT or GFP-CPAP/MT and HA-HBx using anti-HA antibody. The immunoprecipitates were further detected by Western blot analysis by antibodies as indicated. The interaction ability between GFP-CPAP and HA-HBx is indicated as ratio. **(D)** The interaction between CPAP and HBx was determined by *in situ* PLA using anti-CPAP and anti-GFP antibodies (CPAP+, GFP+). The red spots represent interacting complexes of CPAP and HBx. Cells stained with anti-GFP antibody only (CPAP-, GFP+) were used as a negative control. The nuclei were stained with DAPI (blue). **(E)** NF-κB-responsive transcriptional activity (left) and *IL8* promoter activity (right) in cells with GFP-CPAP/WT or GFP-CPAP/MT combined with or without HA-HBx were determined by reporter assay. Error bars represent the mean ± SD of three independent experiments, each performed in triplicate (*, *p* < 0.05; **, *p* < 0.01).

**Figure 3.**
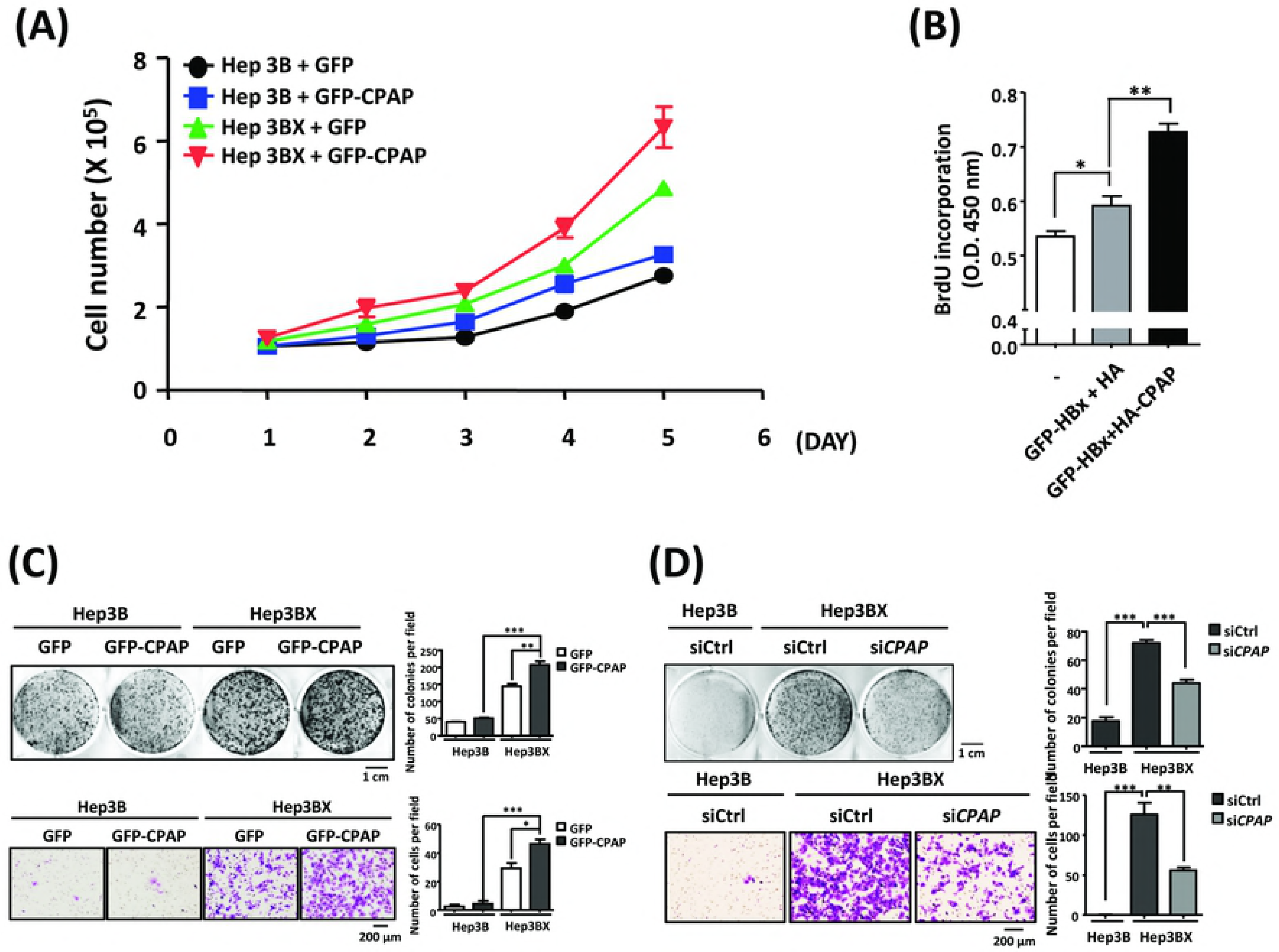
CPAP expression increases HBx-induced cell growth and proliferation of HCC cells. (A) Proliferation of Hep3B and Hep3BX cells with ectopic expressed GFP or GFP-CPAP were determined by counting cell numbers every 24 h under monolayer culture conditions. **(B)** Cell proliferation was examined by BrdU incorporation assay in Hep3B cells with GFP-HBx/HA or GFP-HBx/HA-CPAP expression. The mean ± SD was obtained from three independent experiments (*, *p* < 0.05; **, *p* < 0.01). **(C)** Hep3B and Hep3BX cells with GFP or GFP-CPAP expression were subjected to colony-formation assay (upper) and *in vitro* trans-well migration assay (lower). **(D)** Hep3BX cells with *CPAP* siRNA (si*CPAP*) or control siRNA (siCtrl) were plated for colony-formation assay (upper) and *in vitro* trans-well migration assay (lower) as mentioned above (Figure 3C). Hep3B with control siRNA (siCtrl) was used as a control. The number of cells was counted in 5 randomly selected fields. Data presents as the mean ± SEM in three independent experiments (*, *p* < 0.05; **, *p* < 0.01; ***, *p* < 0.001).

**Figure 4.**
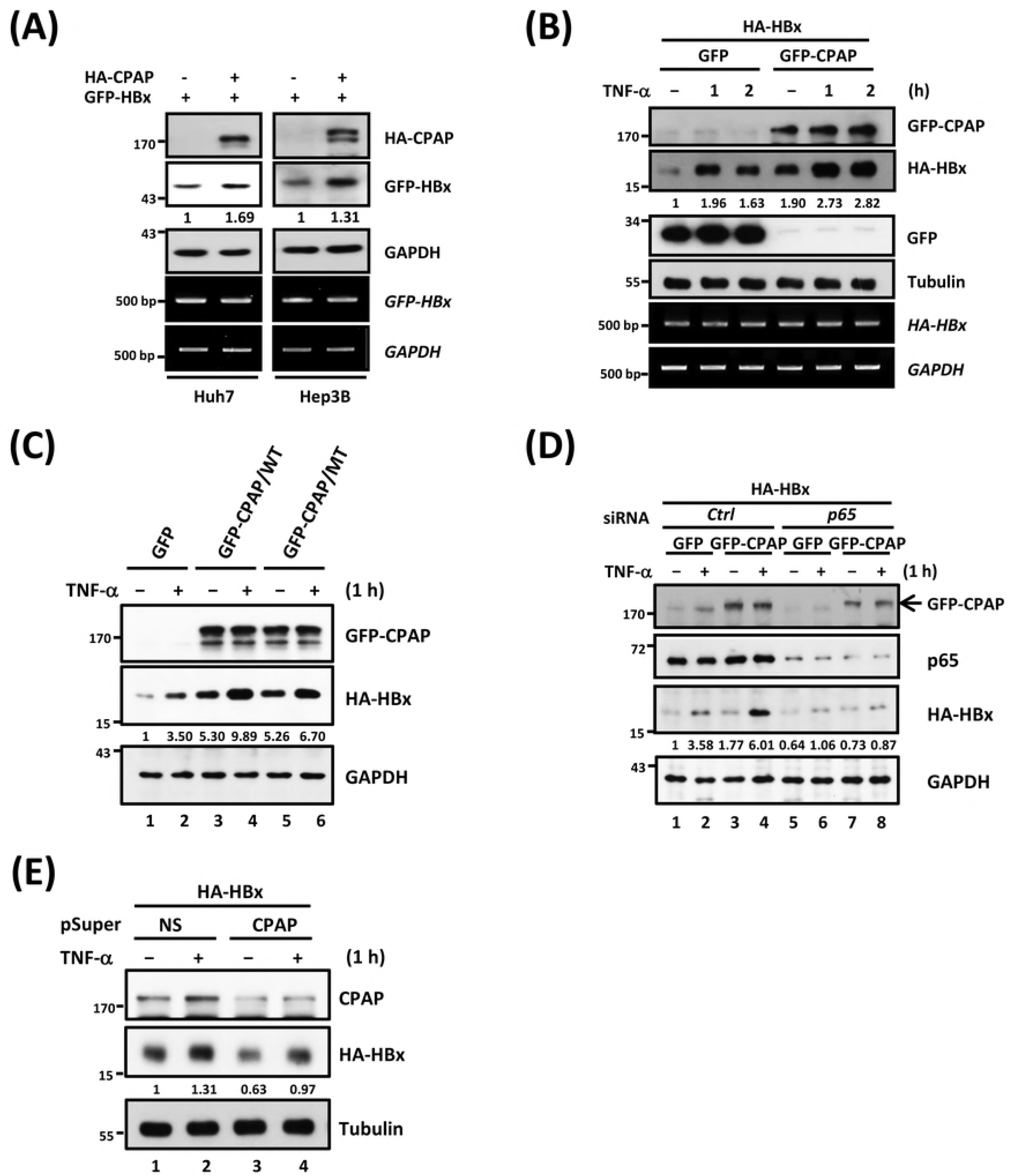
CPAP augments TNF-α-mediated HBx stabilization in a NF-κB-dependent manner. (A) HCC cells with HA-CPAP or GFP-HBx were analyzed by Western blot analysis and RT-PCR. **(B)** Huh7 cells with HA-HBx and GFP or GFP-CPAP were treated with TNF-α (10 ng/ml) for 0, 1, and 2 h. The expression level of GFP, GFP-CPAP, and HA-HBx was determined by Western blot and/or RT-PCR as described in Fig. 1A. **(C)** Western blot analysis of Huh7 cells with GFP-CPAP/WT expression increased HA-HBx stability upon TNF-α treatment (10 ng/ml, 1 h) compared with GFP-CPAP/MT or GFP control. **(D)** Huh7 cells were transfected with non-targeting siRNA (*Ctrl*) or *NF-κB/p65* siRNA (*p65*) for 24 h. The protein levels were examined by Western blot analysis after further transfection of HA-HBx and GFP or GFP-CPAP for an additional 24 h with TNF-α treatment for 1 h.Huh7 cells were transiently transfected with HA-HBx and pSuper/CPAP or.pSuper/NS control for 24 h. Protein expression of CPAP and HA-HBx was determined after treating cells with TNF-α for 1 h. The expression levels of HA-HBx are indicated as ratio, and three independent experiments were performed.

**Figure 5.**
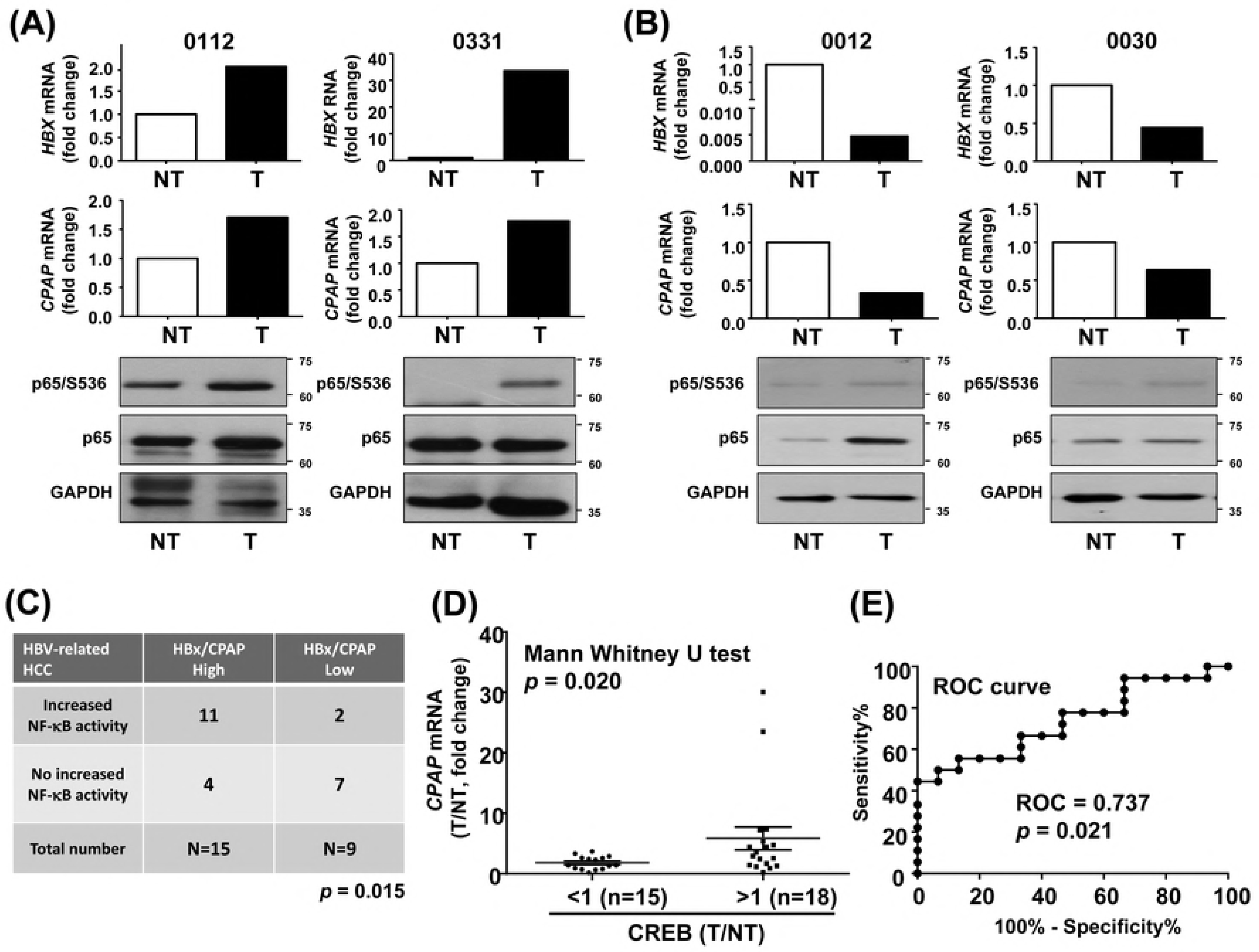
Augmented NF-κB activity in CPAP-overexpressed HBx positive HCC. (A-B) HCC tissues with (A) or without (B) overexpressed CPAP and HBx were collected for Western blot analysis to detect the expression level (p65) and the activation status (p65/S536) of NF-κB. GAPDH was used as a loading control. Expression levels of *CPAP* and *HBx* mRNAs in HCC were detected by Q-PCR, normalized by *actin* mRNA (NT, non-tumor part; T, tumor part). Total of twenty-four HCC tissues with (N=15) or without (N=9) overexpressed HBx/CPAP were analyzed, and two representative cases for both (A) and (B) are shown. **(C)** The correlation between CPAP/HBx expression and activated NF-κB was statistically significant by Chi-square analysis; p=0.015. **(D)** The expression level of *CREB* mRNA is determined in HBx-positive HCC by Q-PCR. A total of thirty-three specimens were enrolled, and the correlation between overexpressed CREB (T/NT >1) and CPAP expression was analyzed by Mann Whitney U test (two-tailed); *p*=0.020. **(E)** ROC (receiver operating characteristic curve) analysis of overexpressed CREB and CPAP in HBx-positive HCC from (D) (*p*=0.021).

We further investigated whether the transcriptional up-regulation of *CPAP* by HBx was due to HBx interacting with CREB and binding to the *CPAP* promoter.ChIP assay confirmed that overexpressed CREB can enhance the association of HBx with the *CPAP* promoter, re-Chip assay further demonstrated that CREB can form a complex with HBx to bind to the *CPAP* promoter (Figure 1E). By contrast, CREB and HBx did not associate with the *CPAP*/M1 promoter (Figure 1F). These results indicated that the *CPAP* promoter -86 to -79 bp is a *cis*-regulatory element for HBx-mediated transcriptional activation; HBx enhances *CPAP* expression by binding onto the *CPAP* promoter through the association with CREB.

### CPAP and HBx cooperatively enhance NF-κB activation

It has been well recognized that NF-κB is a transcription factor regulated by HBx, and contributes to tumorigenesis by activating the expression of several genes involved in the immune response, inflammation, cell proliferation and the prevention of apoptosis(32). On the other hand, it has been reported that CPAP is essential for the transcriptional activation of NF-κB and synergistically increases HBx-mediated NF-κB activation(22, 29). Considering the data from clinical specimens of HBV-HCCs and the critical role of CPAP in NF-κB activation, we investigated the molecular functions of CPAP and HBx in NF-κB signaling. The results showed that the either overexpression of CPAP or HBx both increases the transcriptional activity of NF-κB, and the co-expression of CPAP and HBx induced a substantial increase in NF-κB-responsive transcriptional activity (Supplementary Figures 6A, left panel). Knockdown of *CPAP* by siRNA diminished the HBx-induced NF-κB activation (Supplementary Figures 6A, right panel), confirming that CPAP is essential for HBx-enhanced NF-κB activation. The same effects were found using the *IL-8* promoter, activation of which is dependent on NF-κB activity (Supplementary Figures 6B and 6C). Individual expression or co-expression of CPAP and HBx increased the expression of the NF-κB downstream targets *TNF-α, IL-8* and *ICAM-1* (Figure 2A), whereas *CPAP* knockdown reduced the induction of NF-κB-mediated gene expression by HBx (Figure 2B).

**Figure 6.**
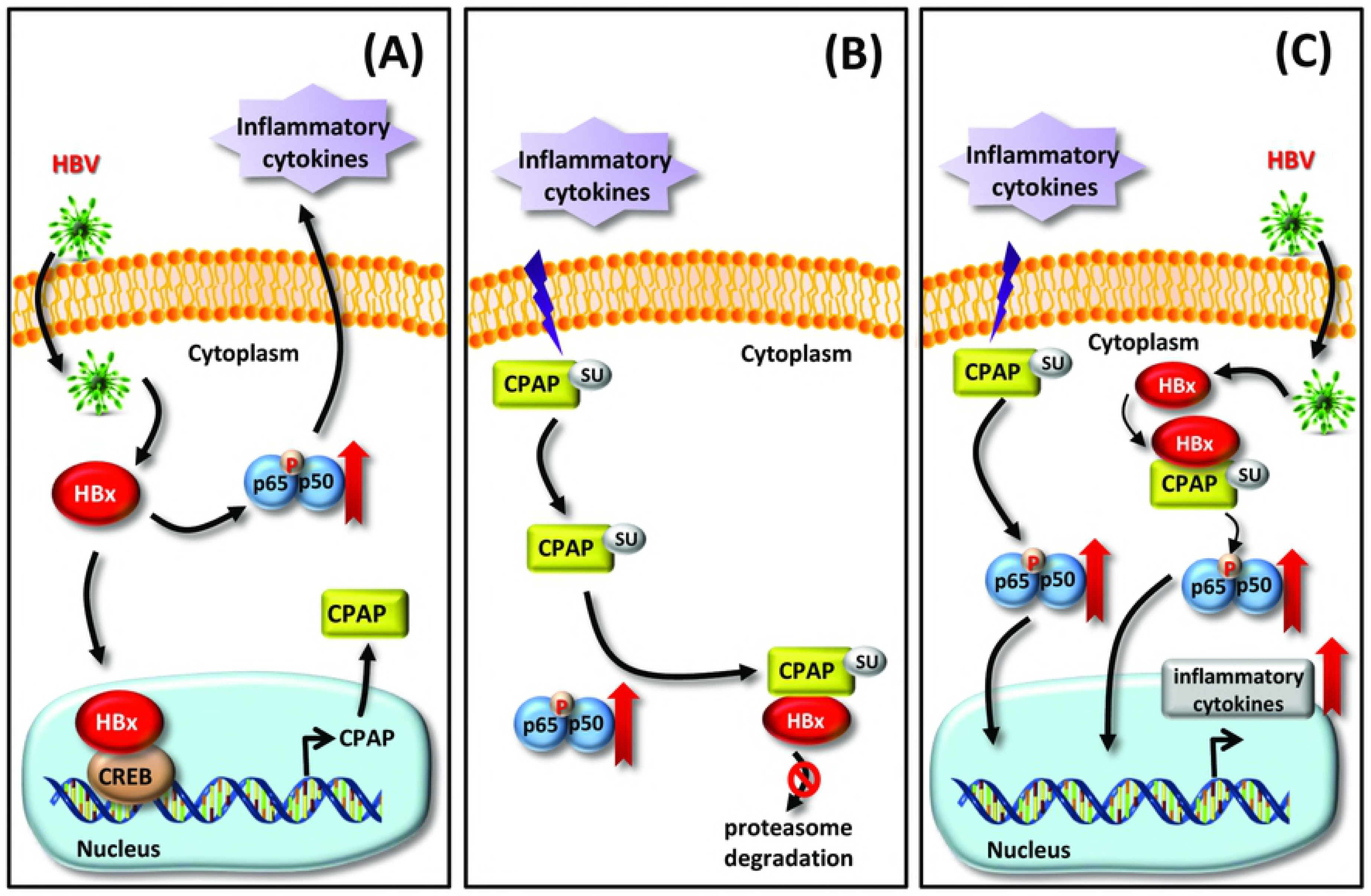
Model of the CPAP/HBx interaction in hepatocarcinogenesis. HBV infection triggers immune responses and later develops into chronic liver diseases, such as hepatitis, fibrosis or cirrhosis. The further production of inflammatory cytokines induced by HBx can modulate the SUMO modification of CPAP and subsequently enhance IKK-mediated NF-κB activation. **(A)** In this study, we demonstrate that HBx associates with the transcription factor CREB to up-regulate *CPAP* transcription. HBx promotes NF-κB activity to enhance the production of inflammatory cytokines. **(B)** Meanwhile, inflammatory cytokines increases HBx stabilization, which is enhanced by CPAP expression and dependent on NF-κB activity. **(C)** CPAP interacts with HBx to further promote HBx-induced NF-κB activation and hepatocellular transformation, finally resulting in hepatocarcinogenesis.

To further investigate how CPAP and HBx cooperate to enhance NF-κB activity, we first checked the association between CPAP and HBx in the NF-κB pathway. Co-immunoprecipitation analysis and *in situ* PLA indicated that CPAP interacts with HBx (Figures 2C, 2D and Supplementary Figure 7A). Our previous report indicated that TNF-α-mediated SUMO-1 modification of CPAP is important for NF-κB activation(22). Here, we investigated whether SUMO modification of CPAP is required for HBx-induced NF-κB activation. The results showed that SUMO-deficient CPAP did not interaction with HBx (Figures 2C, 2D and Supplementary Figure 7A), and decreased the HBx-enhanced NF-κB transcriptional activity (Figure 2E). Furthermore, the interaction between CPAP and HBx was increased upon TNF-α stimulation (Supplementary Figure 7B). These findings suggest that SUMO modification is essential for CPAP to associate with HBx and cooperatively enhance NF-κB activity.

### SUMO modification is required for CPAP to promote HCC proliferation and tumorigenicity

Next, we examined the role of CPAP in HCC development. HepG2 cells that stably expressed GFP-CPAP/WT exhibited a higher proliferative rate and increased colony formation compared with GFP-CPAP/MT and control cells (Supplementary Figure 8A). Moreover, *NF-κB/p65* knockdown decreases CPAP-enhanced colony formation in Hep3B cells (Supplementary Figure 9), suggesting that the CPAP-enhanced proliferation of HCC is dependent on NF-κB activity. To further determine the effect of CPAP on hepatoma cell growth *in vivo*, GFP-CPAP/WT or GFP-CPAP/DM stably expressed HCC cells were subcutaneously injected into NOD/SCID mice. The results showed that overexpression of CPAP/WT significantly increased tumor growth compared with CPAP/DM (Supplementary Figure 8B and Supplementary Figure 10). These results suggest that CPAP promotes HCC growth both *in vitro* and *in vivo*.

### CPAP and HBx cooperatively enhance HCC proliferation, migration and tumourigenesis

Next, we evaluated the contribution of CPAP in HBx-mediated hepatocarcinogenesis by investigating the effects of CPAP on the tumorigenic properties of HCC cells. The results indicated that Hep3B cells expressing either CPAP or HBx showed a higher proliferative rate and colony number than Hep3B cells; co-expression of CPAP significantly increased HBx-enhanced cell growth (Figures 3A, 3B and 3C/upper panel). Moreover, expression of CPAP in Hep3BX cells enhanced HBx-induced cell migration ability (~1.5 fold, Figure 3C/lower panel). To further validate the effect of CPAP in these observations, CPAP was knocked down in Hep3BX cells, and the results showed an impairment of HBx-induced cell growth and migration (Figure 3D). These results support that CPAP is crucial for HBx-induced tumorigenesis in HCC.

### CPAP is required for maintaining TNF-α-mediated HBx protein stability through enhancing the NF-κB activity

Interestingly, we found that the protein expression of GFP-HBx was increased in HA-CPAP-overexpressing cells (Figure 4A), no changes in *GFP-HBx* mRNA expression were observed (Figure 4A), indicating that the prolonged protein stability of GFP-HBx may under the regulation of HA-CPAP. Previous study demonstrated that TNF-α can induce a notable accumulation of HBx by increasing protein stability due to reduced proteasomal degradation through NF-κB signaling(33). Our previous report showed that CPAP is a co-activator of IKK-mediated NF-κB activation in response to TNF-α treatment(22). Therefore, we further clarified whether CPAP can increase the protein stability of HBx through enhancing TNF-α-induced NF-κB activation. The results showed that the highest level of HBx protein was observed 1 h after TNF-α treatment, and then slowly decreased in a time-dependent manner (Supplementary Figure 11A). Overexpressed CPAP significantly augmented HBx protein expression but not mRNA expression in response to TNF-α treatment (Figure 4B). To verify whether the accumulation of HBx protein induced by CPAP is associated with the increased stability of HBx, HCC cells with HA-CPAP and GFP-HBx expression were treated with the protein synthesis inhibitor cycloheximide, and the level of GFP-HBx in the absence or presence of TNF-α was determined at various time points. The results showed that CPAP significantly increased TNF-α-induced HBx stabilization (Supplementary Figure 11B).

Because SUMOylated CPAP is important for TNF-α-mediated NF-κB activation(22), therefore, we investigated whether SUMOylation of CPAP is necessary for HBx stabilization. The results showed that CPAP/WT increased HBx protein expression, whereas the CPAP SUMO-deficient mutant had only a minor effect on maintaining HBx protein expression even in the presence of TNF-α (Figure 4C, compare lanes 4 and 6). According to our previous study, the SUMO-deficient CPAP mutant impaired TNF-α-induced NF-κB activation(22), suggesting that CPAP may be involved in TNF-α-induced HBx stabilization through increased NF-κB activity. To test this idea, we knocked down *NF-κB/p65* and evaluated the effect of CPAP-enhanced TNF-α-induced HBx stabilization. The results showed that the expression of HA-HBx was markedly reduced in *NF-κB/p65* knocked down cells even in the presence of TNF-α (Figure 4D, compare lanes 2 and 6). Moreover, the overexpression of CPAP could not enhance TNF-α-induced HBx stabilization in *p65*-knocked down cells (Figure 4D, compare lanes 4 and 8). In addition, after the knockdown of *CPAP*, HBx expression was decreased regardless of TNF-α treatment (Figure 4E, compare lanes 2 and 4). These results suggest that CPAP is required for TNF-α-induced HBx stabilization.

### Overexpressed CREB/CPAP positively correlates with a poor survival rate in HBx-positive HCC

In order to give insight into the clinical impact of the interaction between HBx and CPAP, we checked the activation status of NF-κB in CPAP-overexpressing HBx-positive HCC. The result indicated that HCC tissues with overexpressed HBx and CPAP has an increased activation of NF-κB (Figure 5A), whereas HCC tissues without HBx and CPAP overexpression present no enhanced NF-κB activity (Figure 5B). The correlation between enhanced NF-κB phosphorylation with overexpressed HBx and CPAP is statistically significant (Figure 5C). Overexpressed CREB is positively correlated with an augmented CPAP level in HBx-positive HCC (Figures 5D, 5E). Importantly, overexpressed CPAP/CREB showed a poor disease-survival rate than instances with a lower expression level of CPAP/CREB (Supplementary Figure 12A). We also analyzed the TCGA data set to check the overall survival rate using SurvExpress biomarker validation tool(34), and the results indicated that an increased expression of *CREB* and *CPAP* mRNAs exists in the high-risk group with a lower survival rate (Supplementary Figure 12B).

## Discussion

HBV integrates its viral DNA into the host cellular DNA to disrupt or promote the expression of cellular genes which then influences cell growth and differentiation. HBV infection is a crucial factor in chronic or acute hepatitis and contributes to the development of HCC(13). Our study proposes a model of the relationship between CPAP and HBx in HCC development. Chronic HBV infection in liver cells triggers host immune responses and results in hepatitis, and the release of inflammatory cytokines can induce SUMO modification of CPAP, as previously shown(22). HBx transcriptionally up-regulates the expression of *CPAP* by interacting with CREB (Figure 6A). Meanwhile, SUMOylated CPAP further enhances NF-κB activation and inflammatory cytokine expression; and SUMO modification of CPAP leads to its further association with HBx and increases TNF-α-induced HBx protein stabilization in an NF-κB/p65-dependent manner (Figure 6B). The reciprocal regulation of CPAP and HBx cooperatively increases NF-κB activity and finally contributes to hepatocarcinogenesis (Figure 6C).

Previous studies have shown that HBx is expressed in human liver specimens with high necroinflammatory activity(35). The chemokine-dependent recruitment of inflammatory cells by HBx involves the activated NF-κB pathway. The NF-κB-induced expression of cytokines, including interleukin-1 (IL-1), IL-6, TNF-α,and interferon-γ (INF-γ), plays an important role in inflammatory processes depending on the duration of inflammation. Remarkably, IL-6 and TNF-α can activate NF-κB in turn to increase the stability of HBx protein in an NF-κB-dependent manner(33). Therefore, it is postulated that positive feedback between HBx and proinflammatory cytokines produced by NF-κB in the microenvironment in the liver leads to hepatic inflammation in chronic hepatitis B. It does not known whether additional proteins can be recruited to enhance HBx stability. In the present study, we demonstrated that CPAP contributes to the protein stability of HBx through increased NF-κB activity (Figure 4). CPAP has been previously identified as a co-activator of NF-κB and as a scaffold protein for the association of IKKs and NF-κB(22, 29). We propose that CPAP enhances NF-κB activation to increase the expression of inflammatory cytokines, further resulting in TNF-α-induced HBx stabilization. Moreover, if the CPAP/NF-κB pathway could be disrupted, the HBV-infected liver cells might be able to reduce HBx-mediated HCC development. As mentioned in previous studies, C-terminal modification of CPAP with SUMO is stimulated by TNF-α and occurs close to the NF-κB/p65-interacting domain of CPAP(22, 29).SUMO modification is also required for the interaction of CPAP and HBx and for the enhanced effect of CPAP on TNF-α-induced HBx stabilization(22). It remains to be elucidated whether the complex of SUMOylated CPAP and NF-κB/p65 can prevent.the association of HBx with proteasome subunits to increase HBx protein stability.

Here, we demonstrate a novel regulatory mechanism between CPAP and HBx in inflammation-related HCC development. The HBx/inflammatory cytokines/CPAP regulatory loop resulted in marked NF-κB activation in HBV-associated HCC, which provides a microenvironment for tumor development. Additionally, overexpressed CREB/CPAP indicated a poor prognostic value in HBV-associated HCC. Taken together, our findings provide an alternative therapeutic target in the NF-κB pathway to reduce the immunodeficiency caused by NF-κB inhibition, which may lead to a novel therapeutic strategy for HBV-association HCC and other chronic inflammatory diseases.

## Materials and Methods

### Xenograft tumorigenicity assay

Hep3B cells (2×10^6^) stably expressed GFP, GFP-CPAP/WT, GFP-CPAP/MT were injected subcutaneously into the right flank region of 5-week-old male NOD-SCID mice. The tumor volumes were measured using calipers every 3 days. Tumor size was measured using the formula: length X width^2^ X 0.5. At 28 days after injection, all mice were sacrificed and tumors were weighed and photographed.

### Co-immunoprecipitation assay and Western blot analysis

Co-Immunoprecipitation assay and Western blots were performed as described(22). Lysates were analyzed by immunoblott analysis using the specific antibodies as indicated in the text. Specific bands were detected with a horseradish peroxidase-conjugated antibody and revealed by an enhanced chemiluminescence (ECL) Western blot system (PerkinElmer).

### Chromatin immunoprecipitation (ChIP) and re-chromatin immunoprecipitation analysis

The ChIP assay was performed as described previously(36). Briefly, the DNA-protein complex was eluted in elution buffer (1x TE buffer containing 1% SDS) with rotation at room temperature for 15 min, and immune complex crosslinking was reversed by heating to 65 °C overnight, followed by treating with 100 μg/ml of proteinase K at 50 °C for 1 h. DNA was extracted twice with phenol/chloroform and precipitated with ethanol. The DNA pellet was re-dissolved in H_2_O and subjected to Q-PCR amplification using specific primers for the *CPAP* promoter (sequence information provided in Supplementary Table 1). For re-ChIP, chromatin immunoprecipitates from the first ChIP were desalted by re-chip buffer (50 mM Hepes, 140 mM NaCl, 1 mM EDTA, 1% TritonX-100, 0.1% NaDOC, 0.1% SDS, and 0.5mM PMSF), incubated with the second antibody or normal-serum control, and processed as above.

### Cell culture, reagents and antibodies

HCC cell lines Huh7, SK-Hep1, Hep3B, Hep3BX, HepG2, and HepG2X were maintained in Dulbecco’s modified Eagle’s medium (DMEM, Sigma, St. Louis, MO). Hep3BX and HepG2X are HBx stably expressed Hep3B and HepG2 cells. HepAD38/Tet-off cells were grown in DMEM supplemented with 200 mM GlutaMax (Gibco, Carlsbad, CA), 0.3 μg/ml doxycycline, and 400 μg/ml G418. HepG2-hNTCP-C4 cells were cultured with DMEM/F-12 with 200 mM GlutaMax,50 μM hydrocortisone, 5 μg/ml insulin and 400 μg/ml G418. Media was supplemented with 10% fetal bovine serum, 100 μg/ml streptomycin, and 100 units/ml penicillin. The cells were incubated at 37°C in a humidified chamber containing 5% CO_2_ Human TNF-α, cycloheximide and anti-α-tubulin antibodies were purchased from Sigma-Aldrich (St. Louis, MO). The antibody against GFP was purchased from Clontech (Palo Alto, CA). Antibody against HA was purchased from Covance (Berkeley, CA). Antibodies against GAPDH and NF-κB/p65 were purchased from Santa Cruz Biotechnology (Santa Cruz, CA). The ON-TARGET plus SMARTpool siRNA for knockdown of CPAP and NF-κB/p65 were obtained from Dharmacon RNA Technologies (Lafayette, CO). sh*GFP* and sh*CREB* were obtained from National RNAi core facility in Taiwan.

### Inducible expression of HBV and HBV infection

HepAD38/Tet-off cells were grown in DMEM with 0.3 μg/ml doxycycline, and the production of HBV was induced by removing doxycycline. HepG2-hNTCP-C4 cells were seeded at 5×10^5^ cells/well in a 6 cm collagen-coated plate. HBV derived from HepAD38/Tet-off cells was infected into HepG2-NTCP-C4 cells at 1000 genome equivalent (GEq)/cell; after 12 h infection, the media was replaced with fresh DMEM/F12 containing 2% FBS, 4% PEG8000. The infected cells were washed three times with PBS at 0 day post infection (dpi), then incubated in fresh DMEM/F12 supplemented with 10% FBS. The cells were collected at 6 days post infection (dpi).for further analysis.

### Luciferase reporter assays

HCC cells were grown in 24-well plate and transfected with a mixture of different vectors as indicated in the figure legends using PolyJet transfection reagent (SignaGen, Rockville, MD, USA) or Lipofectamine reagent (Invitrogen, Grand Island, NY, USA). Luciferase activities were measured with Briteplus (PerkinElmer, Waltham, MA, USA) after 24 or 48 h transfection. All experiments were repeated at least three independent times in triplicate.

### In situ proximity ligation assay (PLA)

HCC cells were grown on sterile cover slips, and after 24 h transfection with HA-CPAP/WT or MT and GFP-HBx, cells were fixed in 3.7% formaldehyde for 10 min and then *in situ* PLA was performed according to the manufacturer’s instructions (Olink Bioscience, Uppsala, Sweden). Two primary antibodies derived from different species were used to recognize CPAP and GFP. Secondary antibodies were species-specific PLA probes. The interaction of proteins was amplified as distinct bright-red spots and detected using a fluorescence microscope (Personal DV Applied Precision, Issaquah, WA).

### RNA extraction and quantitative real-time PCR

Total RNA was extracted using TRIsure reagent (Bioline, London, UK). cDNA was synthesized with the High capacity cDNA Reverse Transcription Kit (Applied Biosystems, Grand Island, NY, USA). Quantitative real time-PCR assay to detect mRNA expression were conducted using iQ(tm) SYBR^®^ Green Supermix (Bio-Rad, Hercules, CA, USA). *Actin* was used as an internal control. Primers specific for human genes are described in Supplementary Table 1.

### Colony formation assay

For clonogenicity analysis, 3~5×10^3^ cells/well were seeded in 6-well plate and cultured in complete medium for 10~15 days. Colonies were fixed with formaldehyde and stained with 0.1% crystal violet.

### Cell proliferation assay

Cells (3×10^3^ cells/well) were seeded in 96-well plate and maintained in complete medium overnight. Cell proliferation was performed using the CCK-8 and BrdU incorporation at indicated times according to the manufacturer’s instructions.

### In vitro trans-well migration assay

Cells were resuspended in serum-free medium and 400 μl of cell suspension (1.5×10^5^cells) were placed into the upper chamber (Millicell chambers, Millipore). Medium containing 10% FBS were added to the lower chamber. After 20 h of incubation, the cells remaining on upper chamber were removed, and the attached cells on the lower membrane were formaldehyde-fixed and stained with 0.1% crystal violet. The migration rate was quantified by counting the migration cells in five random fields using an inverted microscope.

### The Cancer Genome Atlas (TCGA) data set

Data from the TCGA data set was used to analyze the overall survival curves using SurvExpress biomarker validation tool(34).

### Statistical analysis

Statistical differences were assessed between experimental groups using two-tailed and un-paired Student’s t tests. *, *p*<0.05; **, *p*<0.01; ***, *p*<0.001.

### Ethics Statement

The use of clinical HCC specimens was in accordance with the Declaration of.Helsinki. The HCC specimens were obtained from tissue bank of National Cheng Kung University Hospital and this study was approved by the Institutional Review Board (IRB) of National Cheng Kung University Hospital, Tainan, Taiwan (ER-99-347). The usage of all clinical specimens was anonymous.

All animal studies were according to protocols approved by Laboratory Animal Committee of National Cheng Kung University (Approval No. 103234).

## Acknowledgements

The authors thank Nature Publishing Group language editing for help with English editing.

## Supplementary information of legends

Supplementary Figures

There are twelve supplementary figures in this manuscript.

## Supplementary Figure Legends

**Supplementary Figure 1. HBx increases CPAP promoter activity in HCC cell lines.** Huh7, SK-Hep1, HepG2 and Hep3B cells were transiently transfected with GFP or GFP-HBx with *CPAP* promoter for 24 h prior to a reporter assay. GFP-HBx was detected by Western blot analysis. Error bars represent the mean ± SD of three independent experiments, each performed in triplicate (**, *p* < 0.01; ***, *p* < 0.001).

**Supplementary Figure 2. Expression of CPAP is increased in HBV genome-expressing HCC cells. (A)** HBV genome is induced to express in cells without (-) tetracyclin. Expression of HBV core antigen (HBc) was detected by RT-PCR and Western blot analysis. HBc is an indicator for HBV genome expression.**(B)** Total cell lysates from HepAD38 cells with (+) or without (-) tetracyclin were collected to analyze the expression of CPAP. Triplicated cell lysates were analyzed.

**Supplementary Figure 3. The CREB binding site is crucial for *CPAP* promoter activity.** pGL2-*CPAP*/WT or pGL2-*CPAP*/M1, M2 or M1+2 (for detail, please see Figure 1C) were transiently transfected into Huh7 and Hep3B cells prior to a reporter assay. Data presents as the mean ± SEM in three independent experiments, each performed in triplicate (*, *p* < 0.05; **, *p* < 0.01; N.S., no significance).

**Supplementary Figure 4. CREB is essential for HBx-mediated *CPAP* promoter activity. (A)** Hep3B cells with GFP-CREB or/and HA-HBx expression were used in a reporter assay to determine the *CPAP* promoter activity. Expression of GFP-CREB and HA-HBx was detected by Western blot analysis. **(B-C)** *CPAP* promoter activity in Huh7 cells (B) or Hep3B (C) with HA-HBx or/and wild type GFP-CREB (WT) or GFP-CREB/S133A dominant-negative mutant (DN) was determined by reporter assay.**(D)** pGL2-*CPAP*/WT or pGL2-*CPAP*/M1, M2 or M1+2 were transiently transfected into Hep3B cells with GFP-CREB or HA-HBx, and luciferase activity and Western blot analysis were performed. Data presents as the mean ± SEM in three independent experiments, each performed in triplicate (*, *p* < 0.05; **, *p* < 0.01; ***, *p* < 0.001; N.S., no significance).

**Supplementary Figure 5. Evaluation of *CPAP* promoter activity in sh*CREB* transfected Hep3B cells. (A-B)** Three different sh*CREB* (#1, #2 and #3) were used to knock down the expression of endogenous CREB (A), and *CPAP* promoter activity was determined in sh*CREB* #2 and #3 transfected Hep3B cells. **(C)** pGL2-*CPAP*/WT and HA-HBx were co-transfected into sh*GFP* or sh*CREB* (#2 or #3) knocked-down Hep3B cells and then the reporter assay was performed as described above. The numbers indicate the fold change versus sh*GFP* control. Three independent experiments were performed (**, *p* < 0.01).

**Supplementary Figure 6. CPAP is involved in HBx-induced transcriptional activation of NF-κB.** NF-κB-responsive transcriptional activity (**A**) and *IL-8* promoter activity (**B, C**) were examined in Hep3B (B) or Huh7cells (C) using a reporter assay. Cells were transiently transfected with GFP-HBx and HA-CPAP or CPAP siRNA (si*CPAP*) (A), or GFP-CPAP and HA-HBx or pSuper-CPAP (B, C) prior to a reporter assay. The expression levels of individual proteins were detected by Western blot analysis. pSuper/NS or control siRNA (siCtrl) was the siRNA control. Expression of CPAP and HBx was detected by Western blot analysis. RLA, relative luciferase activity. Data presents as the mean ± SEM in three independent experiments, each performed in triplicate (*, *p* < 0.05; **, *p* < 0.01; ***, *p* < 0.001).

**Supplementary Figure 7. The interaction between CPAP and HBx is increased upon TNF-α treatment. (A)** The interaction between CPAP and HBx was determined by *in situ* proximal ligation assay (PLA). Anti-CPAP and anti-GFP antibodies (CPAP+, GFP+) were used to detect the HA-CPAP/wild type (WT) or HA-CPAP/double mutant (DM, K921.975R) and GFP-HBx in SK-Hep1 cells. The red spots represent interacting complexes of CPAP and HBx. Cells stained with anti-GFP antibody only (CPAP-, GFP+) were used as a negative control. The nuclei were stained with DAPI (blue). Expression of HA-CPAP/WT, HA-CPAP/DM and GFP-HBx was detected by Western blot analysis. **(B)** Huh7 cells transfected with HA-HBx and GFP-CPAP or GFP were treated with TNF-α and then harvested for IP assay using anti-HA antibody. The expression level of HA-HBx and the interaction ability between GFP-CPAP and HA-HBx are indicated as ratio.

**Supplementary Figure 8. CPAP promotes proliferation, colony formation, and tumorigenicity of HCC. (A)** Cell proliferation assay (left), BrdU incorporation (middle) and colony-formation assay (right) were examined in Hep3B cells stably expressed with GFP, GFP-CPAP/WT or GFP-CPAP/DM. The data are presented as the mean ± SEM, and three independent experiments were performed (*, *p* < 0.05; **,*p* < 0.01; ***, *p* < 0.001). **(B)** GFP, GFP-CPAP/WT or GFP-CPAP/DM stably expressed Hep3B cells were injected subcutaneously into the right flank of NOD-SCID mice (n=8). Tumor size and weight were examined. The Ki-67 proliferation index of tumor cells is shown.

**Supplementary Figure 9. NF-κB/p65 is essential for CPAP-mediated colony formation of HCC cells.** Hep3B cells with stably expressed GFP or GFP-CPAP were transfected with *NF-κB/p65* (si*p65*) or control (siCtrl) siRNA, and then performed the colony-formation assay. The mean ± SEM were obtained from three independent experiments (**, *p* < 0.01; ***, *p* < 0.001).

**Supplementary Figure 10. Overexpression of CPAP/WT significantly increased tumor growth in a xenograft animal model.** GFP-CPAP/WT or GFP-CPAP/DM stably expressed HepG2 cells were injected subcutaneously into the right flank of NOD-SCID mice (n=6). Tumor volume **(A)** and weight **(B)** were examined. Tumor volume was measured using the formula: length X width^2^ X 0.5. *, *p* < 0.05; **, *p* < 0.01.

**Supplementary Figure 11. CPAP increases TNF-α-mediated HBx protein stabilization. (A)** TNF-α increases GFP-HBx expression. (Upper) Huh7 cells were transiently transfected with GFP-HBx and treated with TNF-α and then harvested for Western blot analysis at indicated time points. (Lower) Quantification showed GFP-HBx expression in TNF-α-treated cells. **(B)** Huh7 cells co-transfected with GFP-HBx and HA-CPAP or HA control were treated with TNF-α for 1 h and then incubated with proteasome inhibitor cycloheximide (200 μg/ml) for 0.5, 1, or 2 h. The cells were harvested for Western blot analysis. The expression levels of GFP-HBx are indicated as ratio.

**Supplementary Figure 12. Co-overexpression of CPAP and CREB is positively correlated with a poor disease-free survival rate in HBx-positive HCC. (A)** Patients with overexpressed CPAP/CREB (T/NT >1, n=12) HCC tissues have a poor disease-free survival rate compared with HCC patients with lower expression level of CPAP/CREB (T/NT <1, n=4). All of these HCC are HBx-positive. **(B)** (Left) The HCC data set from TCGA (n=361) was split into two maximized risk groups, low-risk (green) and high-risk (red), according their prognostic index. *P*-values correspond to log-rank test and *t*-test for the box plot. (Right) Overexpression of *CPAP* and *CREB* mRNAs exists in the high-risk group with a lower overall survival rate (red).

Supplementary Tables

**Supplementary Table 1. Primers used in PCR analysis.**

**Supplementary Table 2. Clinical parameters of the patients with HBV-related HCC who were included in this study.**

## List of supplementary material online

**Supplementary Figures**

**Supplementary Figure Legends**

**Supplementary Tables**

